# Mircubactin B rescues the lethal effect of cell wall biosynthesis mutations in *Bacillus subtilis*

**DOI:** 10.1101/2022.04.24.489322

**Authors:** Bernhard Kepplinger, Xin Wen, Andrew Robert Tyler, Byung-Yong Kim, James Brown, Peter Banks, Yousef Dashti, Eilidh Sohini Mackenzie, Corinne Wills, Yoshikazu Kawai, Kevin John Waldron, Ellis Nicholas Edward Allenby, Ling Juan Wu, Michael John Hall, Jeff Errington

## Abstract

Growth of most rod-shaped bacteria is accompanied by the insertion of new peptidoglycan into the cylindrical cell wall. This insertion, which helps maintain and determine the shape of the cell, is guided by a complex protein machinery called the rod complex or elongasome. Although most of the proteins in this complex are essential under normal growth conditions, cell viability can be rescued, for reasons that are not understood, by the presence of a high (mM) Mg^2+^ concentration. We screened for natural product compounds that could rescue the growth of mutants affected in rod-complex function. By screening >2,000 extracts from a diverse collection of actinobacteria, we identified a new compound, mirubactin B, related to the known iron siderophore mirubactin A, which rescued growth in the low micromolar range, and this activity was confirmed by synthesising mirubactin B. The compound also displayed toxicity at higher concentrations, and this effect appears related to iron homeostasis. However, several lines of evidence suggest that the mirubactin B rescuing activity is not due simply to iron sequestration. The results demonstrate a novel antibacterial compound and add to growing evidence that bacterial siderophores have a range of activities beyond simple iron sequestration.

## INTRODUCTION

Gram-positive bacteria surround their cell membrane with a thick layer of peptidoglycan (PG), which gives the bacteria their shape and protects them from fluctuations in internal osmotic pressure.^1^ PG synthesis is highly conserved across bacterial taxa and several components of the PG synthetic machinery are hugely important antibiotic targets. The cell envelopes of Gram-positive bacteria usually contain other highly abundant anionic polymers called teichoic acids (TAs), which can be either anchored into the cell membrane (lipoteichoic acids, LTA) or to the peptidoglycan (wall teichoic acids, WTA).^2^ The specific functions of these polymers are not well understood but this certainly includes a role in metal homeostasis^2,3^ and they can be important virulence determinants in pathogens. ^4–6^

Cell wall synthesis is supported by various synthetic systems. In most rod-shaped bacteria cell elongation is governed by various general elements: a “Rod complex”, which includes a glycosyl transferase, RodA, and associated transpeptidases called class B penicillin-binding proteins (bPBPs); autolytic hydrolases such as CwlO and LytE in *Bacillus subtilis*,^7^ which break bonds to allow cell wall expansion; and a topological regulator, the actin-like protein MreB, and its associated factors MreC and MreD. Some TA-synthetic enzymes and PG precursor synthetic proteins are also probably associated with the Rod complex.^8^

In *B. subtilis*, there are three MreB paralogues, which have overlapping functions.^9^ MreB and Mbl (“MreB-like”)^10–13^ are both essential in typical microbiological growth media but, curiously, this growth deficiency can be rescued by addition of high concentrations of magnesium (Mg) to the culture medium.^14^ Intriguingly, growth of mutants that lack one of several other factors in the Rod complex can also be rescued or greatly enhanced by Mg^2+^ supplementation, including *mreC, mreD* and *rodA*.^14–17^ *B. subtilis*, like most other bacteria, has a second PG synthetic machinery based around bifunctional (glycosyl transferase, transpeptidase) class A penicillin-binding proteins (aPBPs).^18–20^ Growth of mutants affected in the major aPBP gene, *ponA*, are also rescued by Mg^2+^.^21^ The molecular basis for this Mg effect remains unclear. Formstone and Errington suggested two potential mechanisms: stiffening of the cell wall or inhibition of autolytic enzymes.^14,22,23^ Dajkovic *et. al.* subsequently presented evidence to exclude mechanical changes in the cell wall, and provided strong evidence for the reduction of autolytic activity.^23^ Wilson and Garner (2021) identified LytE as a likely target for this inhibition.^24^

The TAs are underexplored potential antibiotic targets for Gram-positive pathogens. Compounds active on WTA^25^ or LTA^26^ have been described but, so far, have not yet entered clinical development. The key enzyme in LTA synthesis, LTA synthase (LtaS), is a particularly interesting antibiotic target because it is essential under normal conditions in *Staphylococcus aureus*,^27,28^ but specific inhibitors would not affect Gram-negative nor many other Gram-positive bacteria. Furthermore, the enzyme’s active site is located outside the cytoplasmic membrane, excluding cell permeability and efflux mechanisms as sources of resistance. Lastly there is no equivalent in mammalian cells.

We previously described the discovery of mutations in *ltaS* as potent suppressors of the Mg-dependence of *mbl* mutants in *B. subtilis*.^29^ This suggested that inhibitors of LtaS might also rescue the growth of an *mbl* mutant cultured in the absence of Mg^2+^, providing the basis for a powerful chemical biology screen for LtaS inhibitors. Even if compounds rescuing the growth of an *mbl* mutant did not directly target LtaS they might still be of interest by providing insights into the molecular basis for the Mg-rescue effect and the functions of TAs.

Actinomycetes have been a rich source of bioactive natural products including many diverse drugs, particularly antibiotics.^30^ By screening a collection of actinomycete extracts we were able to identify, isolate and characterise a small molecule, which we named mirubactin B, which efficiently rescues the growth of an *mbl* mutant in the absence of added Mg^2+^. We used bioactivity-guided fractionation to purify the molecule and determined its structure by a combination of NMR and high resolution mass spectrometry. The molecule turns out to be identical to a fragment of a known siderophore antibiotic called mirubactin.^31,32^ To validate the source of the observed bioactivity, mirubactin B was synthesised, and the synthetic molecule was also able to rescue growth of *mbl* and *mreB* mutants of *B. subtilis* but was toxic at higher concentrations, probably due to iron sequestration. The results provide new insights into the intricate connections between metal homeostasis and cell envelope synthesis systems, as well as highlighting how poorly we understand the complex chemical exchanges between microorganisms inhabiting the soil.

## RESULTS

### A screen for natural product compounds that rescue growth of an *mbl* mutant

Previous results showed that disruption of the *mbl* gene is normally lethal on standard culture media but that the mutant can be rescued by the addition of a high concentration (e.g. 20 mM) of Mg^2+^. Cells of the *mbl* mutant cultured in the presence of Mg^2+^ and then diluted into medium with no added Mg^2+^ failed to grow, except after prolonged incubation, at which time it appeared that suppressor mutations emerged that relieve the Mg dependence, as described by Schirner & Errington (2009).^29^ We established a protocol for culturing *mbl*-mutant cells in 96 well plates. Supplementary Figure 1 shows that, under these conditions, addition of 20 mM Mg^2+^ resulted in growth near to that of isogenic wild type cells (*B. subtilis* 168CA).

Lower concentrations of Mg^2+^ (down to 2.5 mM) allowed equivalent growth rates but with an increasingly long delay. We do not understand the basis of this but a similar effect has been described by Pi et al. (2020).^33^ Agar plate crush extracts (see Methods) of strains from a large collection of diverse and well dereplicated actinomycetes were obtained from Demuris Ltd. A total of 2,070 extracts were screened, from organisms belonging to various Genera, including: *Actinomadura, Amycolatopsis, Dactylosporangium, Gordonia, Micromonospora, Norcardia, Rhodococcus, Streptacidiphilus, Streptomyces* and *Streptosporangium*. The collection is known to contain a high proportion of antibiotic producers, with about 25% of organisms capable of producing secondary metabolites that kill *B. subtilis* in simple plug assays. Five of the strain extracts were found reproducibly to enhance the growth of the *mbl* mutant in the absence of added Mg, giving growth rates similar to 2.5 to 10 mM of Mg^2+^ (representative plots shown in Supplementary Figure 1.

### Genome sequencing of the producer strains

To facilitate purification of the active compounds we tested the supernatants of the five positive strains for the *mbl*-rescuing. Of the five strains, only TW 167, MEX267 and MDA8-470 reproducibly produced under these conditions, so we focused on these three strains. As a method to determine what kind of bioactive natural products these organisms might make and whether they were likely to make similar or different compounds, we carried out whole genome sequencing (WGS) by a combination of Minion and Illumina sequencing methods (*accession numbers will be provided on acceptance*). Analyses of the predicted natural product biosynthetic gene clusters in each of the three strains are shown in Supplementary Table 1 through antiSMASH output of the prioritised strains. This analysis revealed that TW 167 and MDA8-470 are closely related strains, whereas MEX267 is evolutionarily distant.

The nearest described type strains were *Streptomyces drozdowiczii* for TW 167 and MDA8-470 and *Nocardia zapadnayensis* for MEX267. We chose MDA8-470 as the focus of further work since in repeated growth experiments it tended to provide bioactivity more consistently than the other strains.

Purification and structure determination of the active compound from MDA8-470

We purified the active component from supernatants of strain MDA8-470 by activity-guided isolation using the *mbl*-rescue bioassay. MDA8-470 was cultured for 76.5 h at 26 L volume in a stirred tank reactor. The culture broth was treated with XAD-16 resin and absorbed compounds were eluted with methanol. Hydrophobic compounds were removed by partitioning between ethyl acetate (EtOAc) and water (pH 4). The active compound stayed in the aqueous phase which was further purified by C18 flash chromatography followed by LH20 size exclusion in water and then freeze dried to give a light brown powder with a molecular ion by high resolution mass spectrometry (HRMS) of *m/z* 427.1815 [M+H]^+^. We designated this previously unidentified molecule mirubactin B.

### Mirubactin B is a fragment of the siderophore antibiotic mirubactin

The molecular structure of mirubactin B was determined by a combination of HRMS and NMR spectroscopy (Supplementary Table 3 and Supplementary Figure 2). HRMS suggested a molecular formula of C_18_H_25_O_8_N_4_, implying eight double bond equivalents in the structure. ^1^H NMR in D_2_O allowed the identification of 18 hydrogens, suggesting the presence of 7 exchangeable protons, whilst ^13^C NMR showed the presence of a benzene ring and four carbonyl-containing functional groups. Further analysis by ^15^N NMR confirmed the presence of four nitrogen atoms, one amine and three amide-like groups. This, combined with additional 2D experiments, allowed us to propose a linear polypeptide-like structure, including both a 2,3-dihydroxybenzoyl and a hydroxamic acid group typical of bacterial siderophores (Figure 1).

**Figure 1.**
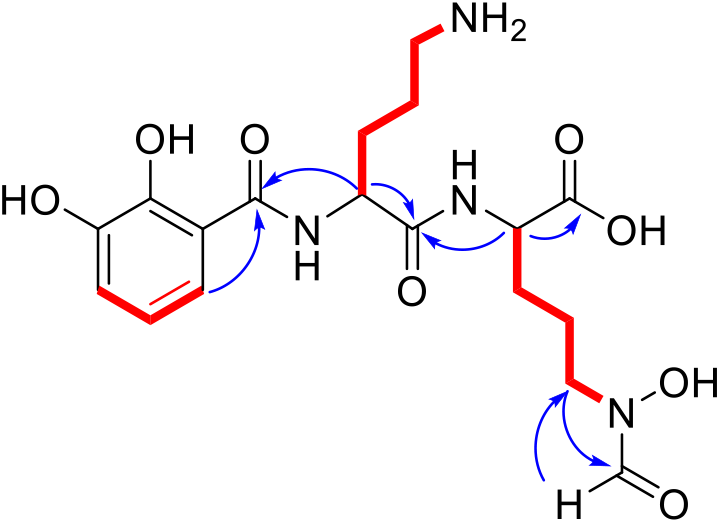
Structural assignment of mirubactin B (COSY = red, selected HMBC = blue)

The structure of mirubactin B turned out to be similar to a family of known compounds including mirubactin (hereafter referred to as mirubactin A), chlorocatechelin A and chlorocatechelin B (Figure 2). In fact, mirubactin B was equivalent to a deletion derivative of mirubactin A, lacking one of its 2,3-dihydroxybenzoic acid moieties. The relationship between mirubactin A and mirubactin B is similar to that of chlorocatechelin A to B^34^.

**Figure 2.**
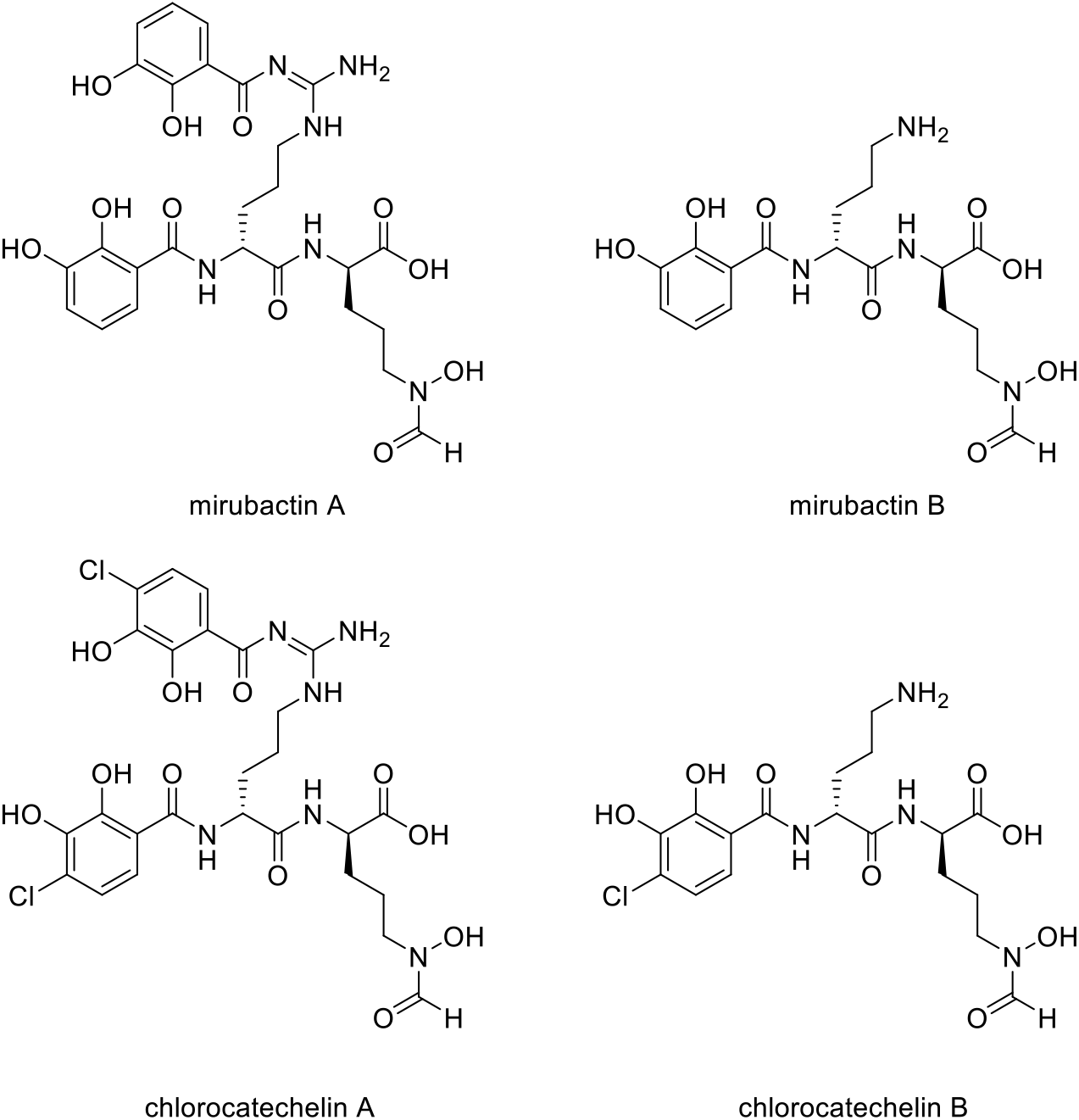
Structures of mirubactin A, mirubactin B, chlorocatechelin A and chlorocatechelin B

Indeed, mirubactin B differs from chlorocatechelin B only in the lack of a chlorine substituent on the aromatic ring. Kishimoto et al. (2015) proposed that chlorocatechelin A spontaneously decomposes to give chlorocatechelin B under acidic conditions, ^34,35^ raising the possibility that mirubactin B has a similar relationship as a degradation product of mirubactin A (Supplementary Figure 4). To check whether mirubactin A was produced by MAD8-470, the strain was cultured in 50 mL of GYM media for three days and, after removing the bacterial cells, the chemical profile of culture media was directly analysed by LC-MS. Mirubactin A was detected in both apo (*m/z* 605.2) and holo (*m/z* 658.1) forms.^32^

Marfy’s analysis was used to determine the stereochemistry of the two constituent ornithine amino acid moieties, which were both found to be in the D-form, matching those of mirubactin A and the chlorocatechelins (Supplementary Figure 3).

We identified a mirubactin-like gene cluster in the genome sequence of MDA8-470 by a combination of antiSMASH analysis and alignments with the mirubactin A gene cluster from *Actinosynnema mirum* DSM 43827 (Figure 3 and Supplementary Figure 2). The gene cluster borders on the gene cluster of coelichelin (an unrelated siderophore) and is consequently not immediately identified as mirubactin by antiSMASH. However, the subcluster analysis found the mirubactin gene cluster with an identity of 72%. The gene cluster appears to be truncated on both sides, while the core biosynthetic pathway is conserved. The biosynthetic genes (*mrbA, mrbC* and *mrbD*; gene designations as assigned by Giessen et al., 2012) for the 2,3-dihydroxybenzoic acid groups (DHAB) appear to be missing from the main cluster. However, genes sharing homology with *mrbC* (57%), *mrbA* (56%) and *mrbD* (55%) were found in an operon approximately 130 kb upstream of the gene cluster. Genes *mrbL-mrbO* appear to be missing in MDA8-470 but they were annotated as unknown function and regulatory in the original mirubactin cluster, and therefore may not be required for the synthesis of mirubactin A.

**Figure 3.**
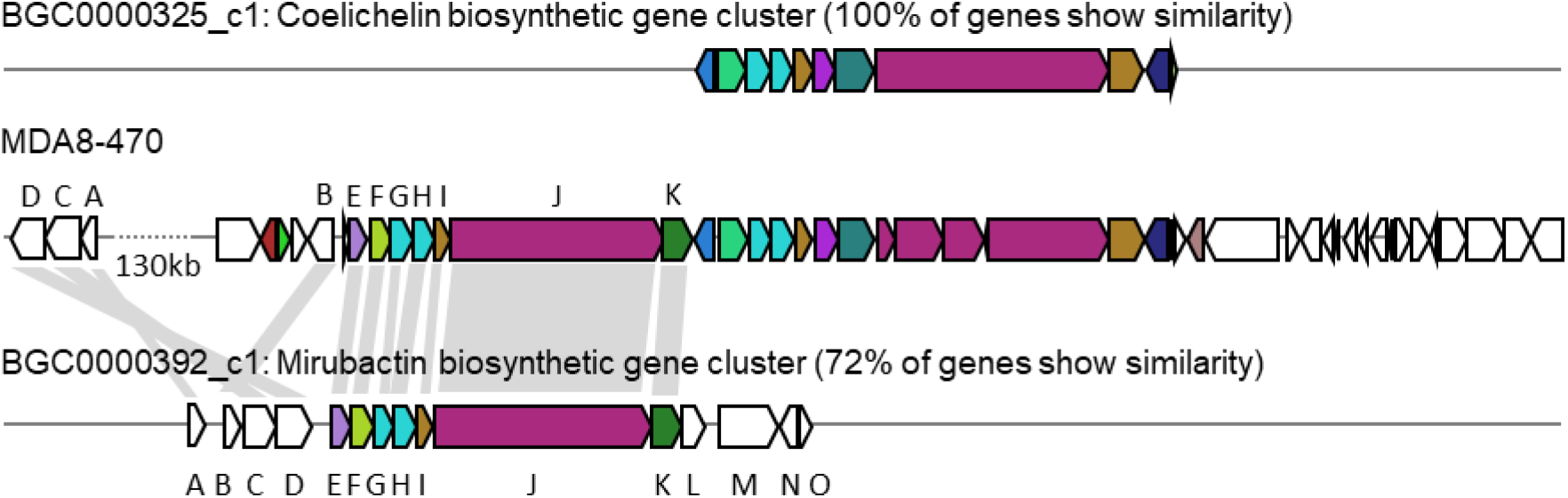
Genetic locus of the mirubactin and coelichelin gene cluster in MDA8-470. Top: coelichelin biosynthetic gene cluster from *Streptomyces coelicolor A3(2)*. Bottom: mirubactin gene cluster from *Actinosynnema mirum* DSM 43827. Homology indicated by grey blocks based on BlastP analysis.

### Synthetic mirubactin B rescues growth of the *mbl* mutant

To confirm that mirubactin B was responsible for the rescue of *mbl* mutant growth we undertook a total synthesis of the compound and compared the activity of this synthetic material to the natural product (Note that details covering the chemical synthesis will be disclosed in a following publication). Synthetic mirubactin B rescued growth at the same concentration as the purified natural compound (Figure 4A & 4B). We therefore conclude that the structure shown in Figure 1 (mirubactin B) is indeed the compound responsible for the *mbl*-growth-rescue effect in extracts from strain MDA8-470. Detailed dose response curves for rescue by the compound (also observed with the original culture extracts) revealed an optimal rescue concentration at 4 µg/mL: above this concentration, growth rescue was progressively less effective (Figure 4A). To ascertain how specific this effect is we also purified mirubactin A and tested it in the *mbl*-recovery assay (Figure 4C). Mirubactin A only showed slight rescue of the *mbl* mutant and required 8 times the concentration (32 µg/mL vs 4 µg/mL). A concentration of 32 µg/mL of mirubactin B already had an adverse effect on the growth of the *mbl* mutant. The failure to rescue growth of the *mbl* mutant at higher concentrations of mirubactin B appeared to be an inhibitory activity, as growth of wild type *B. subtilis* was also impaired at the higher concentrations (Figure 4D). Addition of the optimal concentration of mirubactin B not only rescued growth of the *mbl* mutant, it also largely restored a normal cell morphology, although the cells seemed more prone to chaining than normal.

**Figure 4.**
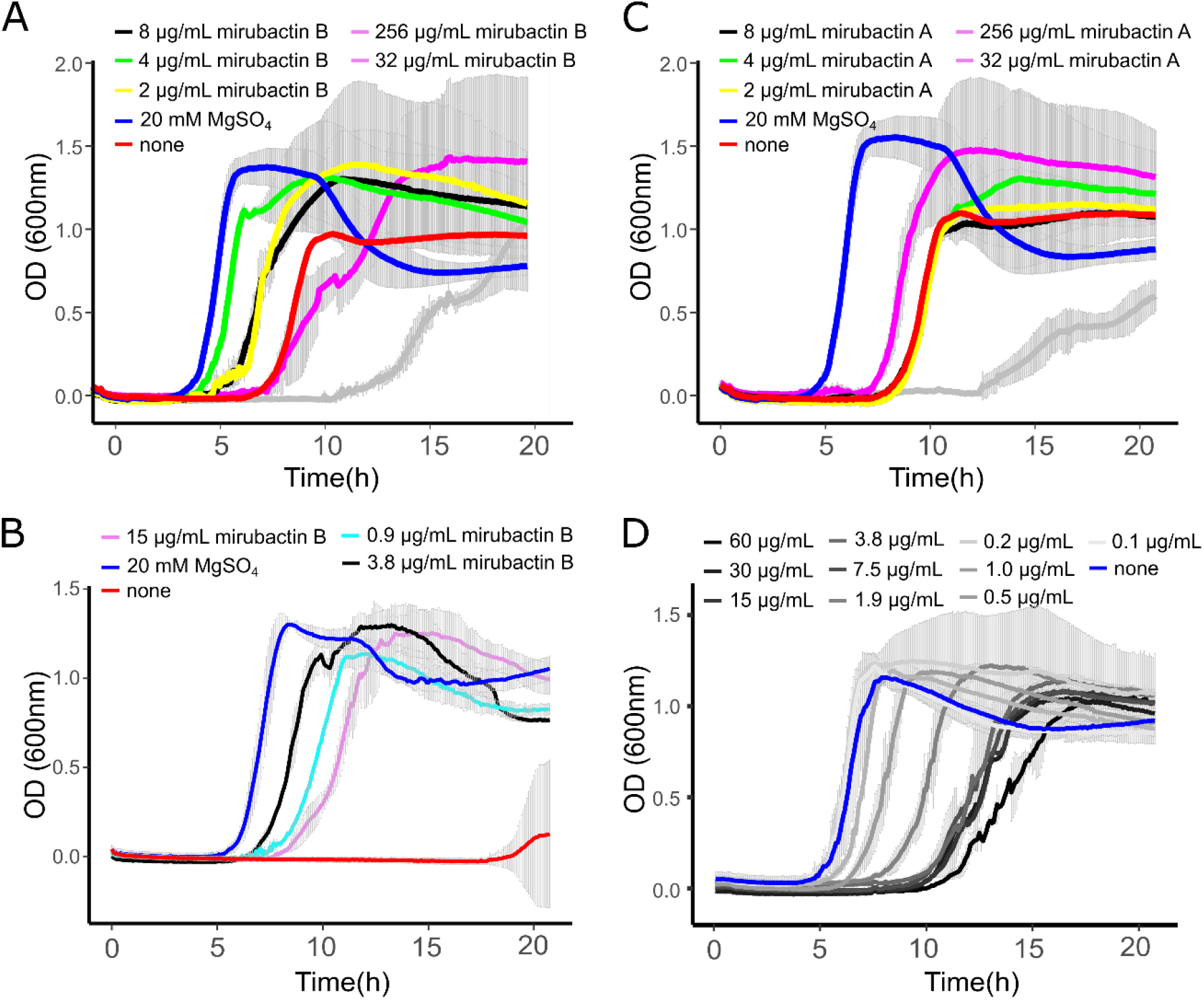
(A) *mbl*-recovery assay with mirubactin B (256 µg/mL grey, 32 µg/mL magenta, 8 µg/mL black, 4 µg/mL green, 2 µg/mL yellow) 20 mM MgSO^4^ (blue), no addition (red). (B) *mbl*-recovery assay with chemically synthesised mirubactin B (15 µg/mL purple, 3.8 µg/mL black, 0.9 µg/mL turquoise), no addition (red), 20 mM MgSO^4^ (blue) (C) *mbl*-recovery assay with mirubactin A (256 µg/mL grey, 32 µg/mL magenta, 8 µg/mL black, 4 µg/mL green, 2 µg/mL yellow), 20 mM MgSO^4^ (blue), no addition (red).(D) Activity of purified mirubactin B against *B. subtilis* 168CA, no addition (blue) and various concentrations of mirubactin B in grey.

### Mirubactin B also rescues the growth of *mreB* but not *ponA* mutants

Previous work has revealed that various mutations affecting PG synthesis are lethal under “normal” culture conditions but can be rescued by addition of high concentrations of Mg^2+^, including *mreB, mreC, mreD, rodA* and *ponA*, and that this is likely due to the inhibition of PG hydrolases (see above). We therefore tested whether mirubactin B could rescue other mutants with a Mg^2+^-dependent phenotype. As shown in Figure 5, mirubactin B rescued the growth of the *ΔmreB* mutant, but not that of the *Δ4* mutant (which lacks *ponA* and the 3 other class A PBPs).^36^ Although growth of the *ΔmreB* mutant was restored, the cells were highly abnormal morphologically, being bloated and lytic (Figure 5).

**Figure 5.**
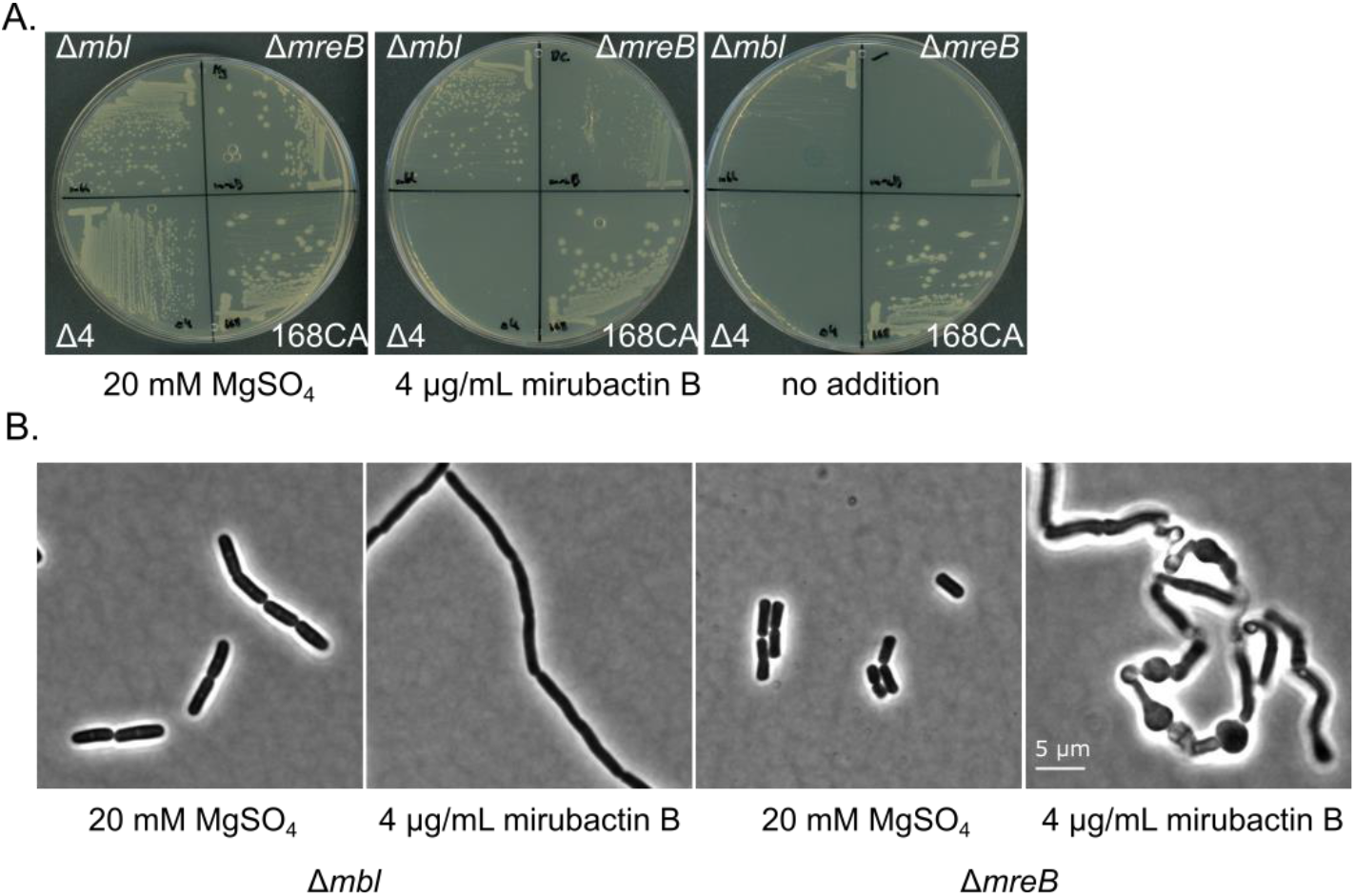
Effects of divalent cations and mirubactin B on several MgSO^4^-dependent cell wall mutants. (A) Growth behaviour of *B. subtilis* mutants Δ*mbl*, Δ*mreB, Δ4* mutant (which lacks *ponA* and genes for the 3 other class A PBPs) on PAB plates in the presence of 20 mM MgSO^4^, 4 µg/mL mirubactin B and in the absence of any additive. (B) Phenotype of *B. subtilis* mutants Δ*mbl* and Δ*mreB* in the presence of MgSO^4^ and 4 µg/mL mirubactin B.

### The inhibitory effect of mirubactin B is associated with iron homeostasis

The mechanism whereby mirubactin B rescues growth of the *mbl* and *mreB* mutants was not clear but we reasoned that a clue might come from further investigation of the inhibitory effect. As one way to investigate this we took a library of *B. subtilis* deletion mutants^37^ and tested the set for growth at a subinhibitory concentration of mirubactin B. The nine most severely affected mutants are listed in Table 1 and the full data set is provided in Supplementary Figure 5. Interestingly five of the nine genes are involved in iron (Fe) homeostasis. These findings suggested that mirubactin B binds iron and that at concentrations above 4 µg/mL, under the conditions used, it causes iron limitation resulting in growth inhibition. To test this idea, we determined the Fe and Mg content of wild type and mutant *B. subtilis* cells by inductively coupled plasma mass spectrometry (ICP-MS) (Supplementary Figure 6) after culture in the presence of Mg or mirubactin B, observing that cells treated with mirubactin B contained significantly lower Fe content. Interestingly, growth-restoring concentrations of mirubactin B did not rectify the Mg deficiency of *mreB* or *mbl* cells, confirming that the metabolite restores their growth through a mechanism distinct from that of Mg^2+^ (Supplementary Figure 6). To confirm the role of Fe, we incubated wild type *B. subtilis* with an inhibitory concentration of mirubactin B (60 µg/mL) and supplemented the culture with FeCl_3_. As shown in Figure 6, growth was partially restored at 3 µM Fe^3+^ and fully restored at 50 µM. This indicates that toxicity of mirubactin B at concentrations above 4 µg/mL is likely due to Fe starvation, consistent with the idea that this compound, like its larger relative mirubactin A, binds iron.

**Table 1.** Deletion strains that exhibit hypersensitivity to mirubactin B

| gene | proposed function according to Subtiwiki<sup>46</sup> |
| --- | --- |
| *feuA* | ABC transporter for the siderophores Fe-enterobactin and Fe-bacillibactin (binding protein), with YusV as ATPase |
| *yusV* | ABC transporter for the siderophores Fe-enterobactin and Fe-bacillibactin, as well as for the siderophores schizokinen and arthrobactin (ATPase) |
| *cpgA* | ribosome assembly GTPase, activity stimulated by ribosomes, may be involved in the maturation of the 30S subunit of the ribosome, detoxification of 4-phosphoerythronate |
| *yqhY* | modulator of lipid biosynthesis |
| *feuB* | ABC transporter for the siderophores Fe-enterobactin and Fe-bacillibactin (integral membrane protein) |
| *feuC* | ABC transporter for the siderophores Fe-enterobactin and Fe-bacillibactin (integral membrane protein) |
| *flhA* | part of the flagellar type III export apparatus (flagellar Type III secretion system), part of the CORE complex required for flagellum and nanotube assembly |
| *fur* | transcription regulator of iron homoeostasis, sensor of Fe sufficiency |
| *sdhA* | succinate dehydrogenase (flavoprotein subunit) |

**Figure 6.**
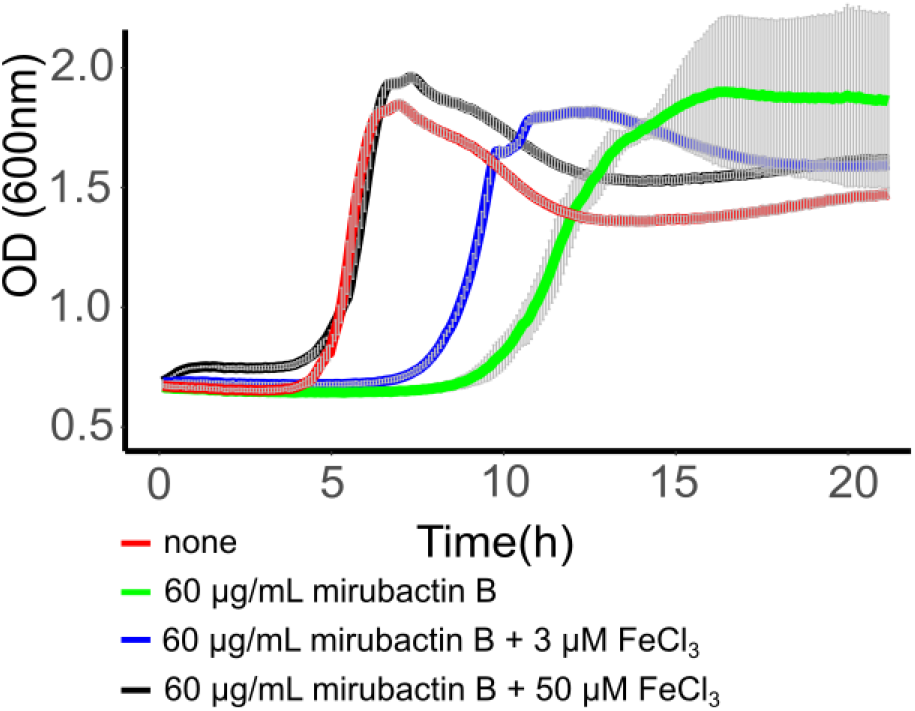
Rescue of mirubactin B toxicity of in the presence of iron. Growth behaviour of *B. subtilis* 168CA; no addition (red), 60 µg/mL mirubactin B (green), 60 µg/mL mirubactin B and 3 µM FeCl_3_ (blue) and 60 µg/mL mirubactin B and 50 µM FeCl_3_ (black)

## DISCUSSION

The main motivation for developing our chemical biology screen was to find potential inhibitors of the LtaS protein, a potential antibiotic target, given that Schirner et al. (2009) had shown that deletion of the *ltaS* gene rescues growth of *mbl* mutants.^29^ However, since Schirner et al. (2009) found suppressor mutations in several other genes, the screen might also identify compounds acting on other targets. Further studies of the hit compounds might also provide insights into the enigmatic ability of Mg^2+^ to rescue the growth of mutants affected in various other cell wall functions. In this work we only screened about 2,000 natural product extracts but this unearthed mirubactin B, plus at least one other structurally and functionally distinct compound that awaits further investigation. It is likely that a screen of larger numbers of extracts would yield yet more interesting compounds, including potential antibiotics. Thus, further screening of synthetic or natural compounds via the *mbl*-rescue assay is warranted.

One of the historical difficulties with screens of natural product compounds was that of dereplicating the hits, i.e. identifying strains likely to be making the same compound and therefore reducing the number of active strains to study via culture scale-up and compound purification. Rapid WGS of the three main hits revealed that two strains were almost identical and therefore likely to be making the same active compound. It was therefore only necessary to follow up one of these strains, illustrating the power of the genomic dereplication approach. The sequencing and analysis also revealed that the third strain is very different to the others and therefore is likely to make an unrelated compound. We intend to follow this up in the future.

The purified active compound turned out to be a smaller version of a known siderophore antibiotic, mirubactin. Total synthesis of the structure deduced for mirubactin B confirmed that this is the active compound in extracts of the producer strain. The genome sequence suggests that strain MDA8-470 contains all necessary genes for biosynthesis of mirubactin and metabolic profile of culture media of strain MDA8-470 confirmed production of mirubactin.. Thus, it seems likely that mirubactin B is either a shunt metabolite in the biosynthesis of mirubactin a or a breakdown product of mirubactin A (Supplementary Figure 4), which would mirror the findings of Kishimoto (2014) with the related chlorocatechelin A and B compounds.^34^ Mirubactin A has a high affinity for Fe^3+^ due to its hexadentate coordination potential based on its DHBA (x2) and hydroxamate moieties. Mirubactin B is expected to have a lower affinity for Fe^3+^ because of the absence of one of the DHBA groups. However, mirubactin B is clearly capable of binding Fe because: first, the inhibitory effect we saw in cultures treated with >=16 µg/ml mirubactin B was exacerbated by various mutations affected in iron uptake or homeostasis; second, the inhibitory effect was alleviated by Fe supplementation in the growth medium. Nevertheless, the rescuing effect of mirubactin B at low Fe concentrations is not simply due to it acting to import iron because we were not able to rescue growth of the *mbl* mutant by addition of mirubactin A. Furthermore, although most actinomycete bacteria produce iron siderophores, only a tiny proportion of the >2,000 culture extracts we tested rescued growth of the mutant. One mechanistic hypothesis worthy of future testing is that mirubactin B moderates siderophore transport in *Bacillus*. This could occur by affecting import through the FeuABC siderophore transporter, in which the FeuA component is known to exhibit promiscuous siderophore binding^38^, by interfering with *feuABC* regulation by interacting with its regulator Btr, which contains a FeuA-like sensory domain^39^, or by modifying the activity of the siderophore esterase, BesA.^40^ Further work will be needed to establish the mechanism whereby mirubactin B rescues growth of the Rod complex mutants. The finding of a natural product compound that has this strange effect on mutants affected in the Rod pathway highlights that there is much still to be learned about the complex chemical genetic interactions between soil microbes.

## METHODS

### Development of a robust high throughput assay for rescue of *mbl-mutant growth*

After exploring several strategies, we fixed on a 96 well format performed in the following manner. A freshly prepared *mbl* mutant strain of *B. subtilis* was obtained by back-crossing DNA from an existing *mbl* mutant into wild type *B. subtilis* 168. The *mbl* mutation was selected on the basis of an associated chloramphenicol or spectinomycin resistance marker, in the presence of 20 mM Mg^2+^, which enables strong growth of the mutant strain. Generation of a fresh *mbl* mutant was necessary to minimise the possibility of emergency and accumulation of growth enhancing suppressor mutations, which could occur when *mbl* mutants were grown at low Mg^2+^ concentrations.

The *mbl* mutant was cultured in nutrient broth (NB) containing 20 mM Mg^2+^ under defined conditions and then diluted 10^−4^-fold into fresh medium containing no added Mg^2+^ and chloramphenicol or spectinomycin (to inhibit growth of the actinomycetes; see below). The mutant culture was dispensed into the wells of a 96 well microtiter plate containing various controls and test samples. To calibrate the assay, several defined concentrations of Mg^2+^ were included. The plates were cultured at 37 °C for 15 hours, and growth was monitored by following the optical density (OD_600_). A wild type culture was also included, with no added Mg^2+^. No growth of the *mbl* mutant was seen in the absence of Mg^2+^ but addition of different concentrations of Mg^2+^ resulted in growth after a delay, which was roughly proportional to the amount of Mg^2+^ added. A typical result is shown below (Supplementary Fig. 1), with time (min) along the x-axis and optical density (OD_600_) on the y-axis. Remaining wells in the plate were occupied by test extracts from the Demuris actinomycete collection. After testing various methods, a freeze-thaw-centrifuge procedure was adopted for preparation of extracts. Individual actinomycete strains were cultured on an appropriate agar growth medium. The agar was harvested and fragmented into 50 mL conical tubes, which were frozen at -20 °C overnight, then thawed and centrifuged. The supernatant was collected, filtered and stored at - 20 °C until used. Positives in the primary assay were identified by growth stimulation similar to Mg^2+^ addition. Candidate extracts were retested using new extracts prepared via regrowth of the actinomycete strains. Five positive producers were identified, strains TW 167, A51P1, Mex267, MDA8-470 and MDA8-566.

### Genome Sequencing and Analyses

The strains TW 167 and MEX267 were grown in 10 mL of liquid GYM and incubated for 3 days at 30°C and 120 rpm. Genomic DNA was extracted using the Quick-DNA HMW MagBead Kit (Zymo Research, cat. no. D6060). Aliquots (300 μL) of the cultured cells were spun at 13,000 × *g* for 2 min to pellet. The cells were resuspended in 200 μL DNA/RNA Shield (Zymo Research, cat. no. R1100-50). Microbial lysis and DNA extraction (including RNAse treatment) were performed according to the manufacturer’s instructions. DNA quality was assessed by agarose gel electrophoresis to ensure no trace amounts of RNA. DNA concentration was assessed using the dsDNA assay on a Qubit fluorometer.

The strains TW 167 and MEX267 were part of a multiplexed nanopore MinION sequencing (12 strains in total) using the Native Barcoding Expansion 1-12 (EXP-NDB104), in conjunction with the Ligation Sequencing Kit (SQK-LSK109). A DNA fragmentation step was not performed. Priming and loading the SpotON flow cell was performed according to the manufacturer’s instructions using R9.4.1 flow cells (FLO-MIN106D, ONT). The flow cell was mounted on a MinION Mk1B device (ONT) for sequencing and was controlled using Oxford Nanopore Technologies MinKNOW software. Base calling and data conversion was performed in parallel using Albacore v1.2.4 (Oxford Nanopore Technologies).The sequences were exported in FASTQ format and used for an assembly with CANU v1.5.^41^

For the Illumina Sequencing the genomic DNA libraries were prepared using the Nextera XT Library Prep Kit (Illumina, San Diego, USA) following the manufacturer’s protocol with the following modifications: input DNA was increased 2-fold, and PCR elongation time was increased to 45 s. DNA quantification and library preparation were carried out on a Hamilton Microlab STAR automated liquid handling system (Hamilton Bonaduz AG, Switzerland). Pooled libraries were quantified using the Kapa Biosystems Library Quantification Kit for Illumina. Libraries were sequenced using Illumina sequencers (HiSeq/NovaSeq) using a 250 bp paired-end protocol.

The resulting reads were mapped onto the contigs produced by Canu using Minimap and the resulting consensus sequences exported. The contigs were annotated using Prokka 1.11.^42^ The corrected genome sequences were analysed with the online version of antiSMASH 6.0.1.^43^ MDA8-470 was sequenced using the Pacific Biosciences single-molecule, real-time DNA sequencing technology performed by the University of Maryland School of Medicine Genomics Resource Centre who also assembled the genome. The resulting four contigs were further polished using paired-end Illumina MiSeq data and annotated using RAST.^44^

### Purification of mirubactin B

The *Streptomyces* isolate MDA8-470 was grown in GYM medium (4 g/L glucose, 4 g/L yeast extract, 10 g/L malt extract) for 76.5h in two stirred tank bioreactors at a combined volume of 26 L. The combined culture supernatant was subjected to a batch absorption of 1.5 kg Amberlite XAD-16N resin. The resin was eluted with 2.5 L of methanol and evaporated to yield in 900 mL of aqueous extract. The extract was twice extracted at pH 4 with an equal volume of ethyl acetate. Bioactivity-guided fractionation showed that the compound stayed in the aqueous phase. We therefore subjected the active aqueous extract in five batches of about 180 mL to 60 g reverse phase flash chromatography from water (0.1 % formic acid) to methanol (0.1 % formic acid) (Gradient 1 column volume (CV) water, 10 CV 0 % - 20 % methanol, 2 CV 20 %-100 % methanol, 1.5 CV 100 % methanol). Mirubactin B eluted at about 10 % methanol. The active fractions were combined and freeze-dried to a viscous oil of 5mL. To remove the formic acid and other minor components we finally subjected the semi pure mirubactin B to a LH20 size exclusion column (85 cm x 3 cm) at a flow rate of 1 mL/min. The active fractions were freeze-dried to give a light brown powder.

### Purification of mirubactin A

A mirubactin producing strain 2161 was cultured in 1 L GYM media for three days. The culture was centrifuged and the supernatant was filtered to remove the cells. The filtrate was passed through a Hypersep C18 (10 g) column using a vacuum pump. The column was then eluted stepwise with 100/0, 90/10, 50/50 and 0/100 of water/methanol. LC-MS analysis showed that mirubactin A eluted in the fraction with 10% methanol. The compound was then further purified on an Agilent 1260 Infinity II preparative HPLC connected to a single-Q mass spectrometer using the following HPLC method: initial isocratic conditions of 5 % acetonitrile for 5 min followed by linear gradient from 5 to 50 % acetonitrile over 45 min; then to 100 % acetonitrile in 1 min; continued by isocratic flow for an additional 9 min at a fellow rate of 12 mL/min. To avoid degradation of mirubactin during chromatography, neat solvents without acid additives were used. A molecular formula of C_26_H_32_N_6_O_11_, consistent with mirubactin A, was generated for the purified compound based on HRESIMS *m/z* 605.2198 [M + H]^+^ (calculated for C_26_ H_33_ N _6_O _11_^+^, 605.2201).

### *B. subtilis* deletion strain library

The strains from a comprehensive *B. subtilis* deletion library^37^ were transferred onto agar plates containing LB agar either with or without 200 μg/ml mirubactin B using a SINGER ROTOR colony pinning robot with 384 long pin pads. The agar plates were incubated overnight at 37°C before being imaged on an S&P robotics SP Imager. The IRIS programme^45^ was used to analyse the agar plate photos and determine individual colony size and the mean of four independent pinning experiments was plotted in Supplementary Figure 5.

### Effect of mirubactin B on the growth of *B. subtilis* 168CA, *Δmbl, ΔmreB* and *ΔponA*

*B. subtilis* wild type 168CA, and isogenic strains bearing mutations *Δmbl, ΔmreB* and *ΔponA* were grown on nutrient agar plates overnight in the presence of 20 mM Mg^2+^. Single colonies of each strain were inoculated into 10 ml PAB medium and grown to an OD_600nm_ of ∼0.1, before diluting 10^−4^ into PAB (Difco Antibiotic Medium 3) containing no Mg^2+^. Mirubactin B was dissolved to 10 mg/mL in water prior to making dilutions in PAB medium to reach the final concentrations shown. FeCl_3_ was dissolved in water to a concentration of 100 mM prior to making dilutions in PAB to reach the final concentrations. Growth was monitored in 96 well microtiter plate. For microscopy analysis the strain was grown to mid logarithmic phase in a 96 well plate prior to microscopic analysis. Microscopy was carried out with Nikon Eclipse Ti (Nikon Plan Apo 1.40 Oil Ph3 objective) and the images were acquired with a Prime 4.2 sCMOS camera (Photometrics) and Metamorph 7 (Molecular Devices).

### Elemental analysis of *B. subtilis* cells

*B. subtilis* strains were cultured in PAB medium, supplemented with 20 mM Mg^2+^ or with 10 g/mL mirubactin B, to OD_600nm_ ∼0.6. Cells were harvested by centrifugation and washed in 1 mL of 20 mM HEPES buffer, pH 7.4, followed by a wash in 20 mM EDTA to remove surface-associated metals, then finally washed twice in PBS to remove trace EDTA. The wet cell weight of the samples was determined and then the pellets were digested in 440 μL concentrated nitric acid before 10-fold dilution for elemental analysis. ICP-MS analysis was performed by Durham University Bio-ICP-MS Facility using a set of matrix-matched standard solutions containing defined concentrations of Mg and Fe, using Sc and Bi as internal standards in all samples and standards.

## Acknowledgements

Work in the Errington lab was supported by a BBSRC Follow on fund grant (BB/FOF/319) and a Wellcome Investigator Award (209500).

Newcastle University for Ph.D. studentships

Dr Alex Charlton (SAGE Mass Spectrometry Facility, Newcastle University) for mass spectrometry support.

Prof William McFarlane (Newcastle University) for NMR support.

Dr Richard Daniel for the use of the *Bacillus subtilis* knock out library.

## Author contributions

**B.K.** investigation, visualization, writing – original draft; **Y.D., X. W., A.R.T., P.B., J.B., E.S.M. & BY.K**. investigation; **C.W. & N.E.E.A.** supervision, **Y.K.**, conceptualization, writing – review & editing; **L.J.W.** supervision, writing – review & editing **K.J.W.** supervision, writing – review & editing, **N.E.E.A., M.J.H.** writing-original draft, supervision. **J.E.** conceptualization, writing – original draft, review and editing, supervision, project administration, funding acquisition.

