## Supplementry Figures and Tables for "Mircubactin B rescues the lethal effect of cell wall biosynthesis mutations in *Bacillus subtilis*"

### Supplementary Info

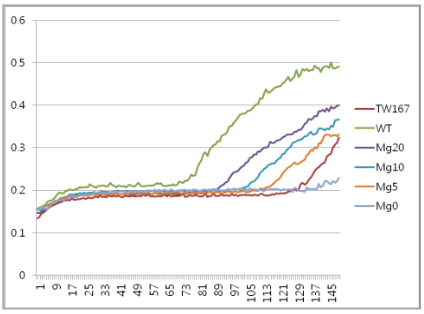


TW 167

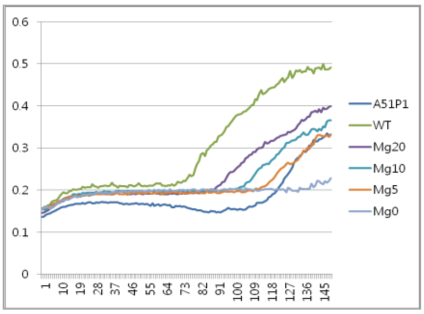


A51P1

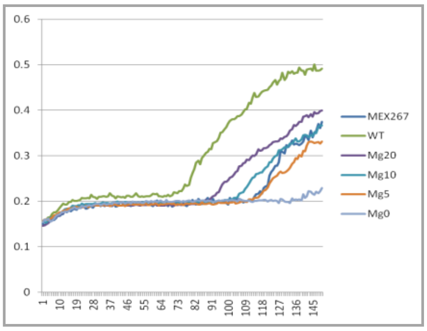


MEX267

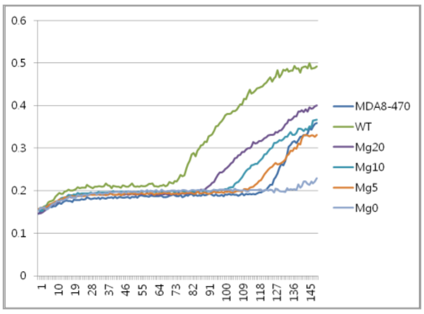


MDA8-470

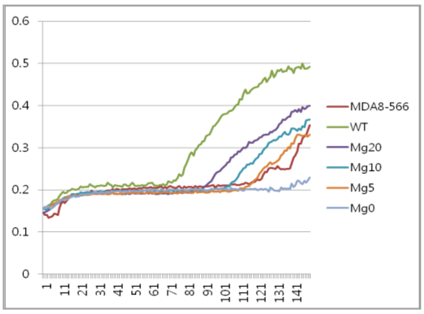


MDA8-566

**Supplementary Figure 1** Confirmation of the *mbl*-rescuing activity of hits from primary extract screening. The *mbl* mutant and wild type control strains were cultured in NB (96 well plates) with added $Mg^{2+}$ as indicated (mM). Fresh agar crush extracts from the indicated strains were added to a concentration of 6% (vol/vol). Growth was recorded as $OD_{600}$.

**Supplementary Table 1** antiSMASH output of the prioritised strains

**MD8A-566**

| Region | Type | From | To | Most similar known cluster | | Similarity |
|---|---|---|---|---|---|---|
| Region 1 | lassopeptide | 49,959 | 72,289 | kanamycin | Saccharide | 1% |
| Region 2 | lanthipeptide-class-iv | 159,332 | 181,995 | venezuelin | RiPP:Lanthipeptide | 100% |
| Region 3 | NRPS,T3PKS | 211,829 | 268,782 | violapyrone B | Polyketide | 28% |
| Region 4 | NRPS | 414,129 | 477,202 | atratumycin | NRP | 7% |
| Region 5 | terpene | 727,150 | 752,475 | hopene | Terpene | 76% |
| Region 6 | RiPP-like | 1,268,156 | 1,279,454 | | | |
| Region 7 | NRPS | 1,493,212 | 1,580,665 | friulimicin A / friulimicin B / friulimicin C / friulimicin D | NRP | 21% |
| Region 8 | siderophore | 1,708,307 | 1,722,357 | ficellomycin | NRP | 3% |
| Region 9 | terpene | 2,087,836 | 2,108,076 | BD-12 | NRP | 17% |
| Region 10 | LAP,thiopeptide | 4,436,710 | 4,471,437 | | | |
| Region 11 | butyrolactone | 4,984,835 | 4,994,482 | | | |
| Region 12 | ectoine | 5,453,618 | 5,463,115 | ectoine | Other | 100% |
| Region 13 | terpene | 5,863,622 | 5,882,692 | steffimycin D | Polyketide:Type II + Saccharide:Hybrid/tailoring | 19% |
| Region 14 | blactam | 6,137,403 | 6,159,229 | tabtoxin | Other | 13% |
| Region 15 | NRPS,betalactone | 6,319,170 | 6,389,723 | coelichelin | NRP | 100% |
| Region 16 | T2PKS,terpene,NRPS | 6,476,466 | 6,602,937 | spore pigment | Polyketide | 83% |
| Region 17 | T2PKS,terpene | 6,620,932 | 6,691,617 | lugdunomycin | Polyketide | 55% |
| Region 18 | T1PKS,PKS-like,NRPS | 6,940,306 | 7,056,880 | salinomycin | Polyketide:Modular type I | 32% |
| Region 19 | T1PKS,butyrolactone,oligosaccharide | 7,056,948 | 7,120,066 | lidamycin | NRP + Polyketide | 39% |
| Region 20 | ectoine,T2PKS,butyrolactone | 7,148,573 | 7,229,947 | kosinostatin | NRP + Polyketide | 22% |

**MEX267**

| Region | Type | From | To | Most similar known cluster | | Similarity |
|---|---|---|---|---|---|---|
| Region 1.1 | T2PKS,butyrolactone,ectoine | 1,882 | 83,259 | kosinostatin | NRP + Polyketide | 22% |
| Region 1.2 | oligosaccharide,T1PKS,butyrolactone,PKS-like,NRPS | 111,705 | 291,764 | C-1027 | Polyketide | 42% |
| **tig00000012** | | | | | | |
| Region 3.1 | T2PKS,terpene | 252,049 | 322,751 | lugdunomycin | Polyketide | 55% |
| Region 3.2 | NRPS,T2PKS,terpene | 343,158 | 467,350 | spore pigment | Polyketide | 83% |
| Region 3.3 | NRPS,betalactone | 553,868 | 624,767 | coelichelin | | |
| Region 3.4 | blactam | 784,515 | 806,353 | tabtoxin | Other | 13% |
| Region 3.5 | terpene | 1,060,336 | 1,081,358 | steffimycin D | Polyketide:Type II + | 19% |

| | | | | | | |
|---|---|---|---|---|---|---|
| Region 3.6 | ectoine | 1,480,132 | 1,490,530 | ectoine | Saccharide:Hybrid/tailoring Other | 100% |
| Region 3.7 | thiopeptide,LAP | 2,471,932 | 2,506,665 | | | |
| tig00000014 | | | | | | |
| Region 5.1 | terpene | 1,176,059 | 1,196,304 | BD-12 | NRP | 17% |
| Region 5.2 | siderophore | 1,561,489 | 1,575,905 | ficellomycin | NRP | 3% |
| Region 5.3 | NRPS | 1,703,576 | 1,791,102 | friulimicin A / friulimicin B / friulimicin C / friulimicin D | NRP | 24% |
| Region 5.4 | RiPP-like | 2,004,590 | 2,015,888 | | | |
| Region 5.5 | terpene | 2,531,096 | 2,557,711 | hopene | Terpene | 76% |
| Region 5.6 | NRPS | 2,817,761 | 2,870,070 | cadaside A / cadaside B | NRP | 14% |
| Region 5.7 | T3PKS,NRPS,NRPS-like | 3,015,453 | 3,072,699 | violapyrone B | Polyketide | 28% |
| Region 5.8 | lanthipeptide-class-iv | 3,102,253 | 3,124,916 | venezuelin | RiPP:Lanthipeptide | 100% |
| Region 5.9 | lassopeptide | 3,211,737 | 3,234,185 | | | |

TW167

| Region | Type | From | To | Most similar known cluster | | Similarity |
|---|---|---|---|---|---|---|
| Region 1.1 | NRPS,terpene | 507,421 | 618,704 | nocobactin NA | NRP + Polyketide | 87% |
| Region 1.2 | NAPAA | 968,144 | 1,002,052 | isatropolone A / isotropolone B / isotropolone C | Polyketide | 9% |
| Region 1.3 | T1PKS | 1,204,735 | 1,249,486 | clorobiocin | Saccharide:Hybrid/tailoring + Other:Aminocoumarin | 7% |
| Region 1.4 | RiPP-like | 1,543,949 | 1,553,780 | | | |
| Region 1.5 | ranthipeptide | 2,000,320 | 2,021,825 | 2'-chloropentostatin / 2'-amino-2'-deoxyadenosine | Other | 6% |
| Region 1.6 | NRPS,RiPP-like | 2,050,975 | 2,168,960 | Sch-47554 / Sch-47555 | | |
| Region 1.7 | NRPS,T1PKS | 3,843,816 | 3,925,042 | pepticinnamin E | NRP + Polyketide | 6% |
| Region 1.8 | NRPS-like | 4,109,167 | 4,152,023 | platensimycin / platencin | Terpene | 5% |
| Region 1.9 | NRPS | 4,348,301 | 4,390,052 | diisonitrile antibiotic SF2768 | NRP | 22% |
| Region 1.10 | T1PKS | 4,441,229 | 4,487,630 | | | |
| Region 1.11 | ectoine | 4,508,666 | 4,519,061 | ectoine | | |
| Region 1.12 | NRPS | 4,595,594 | 4,651,731 | | | |
| Region 1.13 | NRPS | 4,955,436 | 5,019,662 | herboxidiene | Polyketide | 5% |
| Region 1.14 | NRPS | 5,111,873 | 5,170,064 | pentalenolactone | Terpene | 15% |

| | | | | | | |
|---|---|---|---|---|---|---|
| **tig00000002** | | | | | | |
| Region 2.1 | betalactone | 284,078 | 315,945 | | | |
| Region 2.2 | NRPS | 342,406 | 387,295 | heterobactin A / heterobactin S2 | NRP | 63% |
| Region 2.3 | terpene | 645,041 | 664,258 | carotenoid | Terpene | 27% |
| Region 2.4 | T3PKS | 739,330 | 780,499 | | | |
| Region 2.5 | arylpolyene | 803784 | 844956 | echoside A / echoside B / echoside C / echoside D / echoside E | | 11% |
| Region 2.6 | terpene | 1,244,472 | 1,263,989 | | | |
| Region 2.7 | terpene | 1,277,197 | 1,303,501 | A54145 | NRP | 5% |
| Region 2.8 | NRPS-like,NRPS,betalactone | 1,363,396 | 1,437,883 | | | |
| Region 2.9 | terpene | 1,900,050 | 1,918,393 | carotenoid | Terpene | 18% |
| Region 2.10 | terpene | 1,949,854 | 1,970,960 | | | |
| Region 2.11 | betalactone | 2,184,602 | 2,210,492 | | | |
| **tig00000006** | | | | | | |
| Region 6.1 | NAPAA | 101,882 | 135,916 | | | |
| **tig00000007** | | | | | | |
| Region 7.1 | terpene | 133,600 | 154,517 | lavendiol | Polyketide | 6% |
| **tig00000008** | | | | | | |
| Region 8.1 | terpene | 1 | 13,490 | carotenoid | Terpene | 50% |
| **tig00000009** | | | | | | |
| Region 9.1 | NAGGN | 3,031 | 18,158 | bottromycin A2 | RiPP:Bottromycin | 6% |

**Supplementary Table 2** BlastP analysis of the mirubactin gene cluster identified in strain MD8A-470 against the mirubactin gene cluster (BC0000392_c1)

| ID mirubactin cluster | Max Score | Total Score | Query Cover | E value | Per. Ident | Acc. Len |
|---|---|---|---|---|---|---|
| mrbA | 244 | 244 | 96% | 8.00E-86 | 55.89% | 279 |
| mrbB | 296 | 342 | 72% | 5.00E-99 | 57.39% | 399 |
| mrbC | 588 | 588 | 98% | 0 | 57.19% | 553 |
| mrbD | 210 | 210 | 92% | 7.00E-74 | 54.50% | 211 |
| mrbE | 407 | 407 | 89% | 7.00E-147 | 73.74% | 297 |
| mrbF | 426 | 426 | 97% | 2.00E-153 | 69.67% | 334 |
| mrbG | 348 | 348 | 91% | 1.00E-122 | 71.15% | 342 |
| mrbH | 444 | 444 | 88% | 3.00E-160 | 78.07% | 332 |
| mrbI | 417 | 417 | 99% | 6.00E-152 | 77.31% | 262 |
| mrbJ | 3705 | 3705 | 99% | 0 | 62.76% | 3328 |
| mrbK | 566 | 566 | 97% | 0 | 68.05% | 467 |
| mrbL | | | no significant hit | | | |
| mrbM | | | no significant hit | | | |
| mrbN | | | no significant hit | | | |
| mrbO | 30.4 | 30.4 | 28% | 8.00E-04 | 42.86% | 203 |

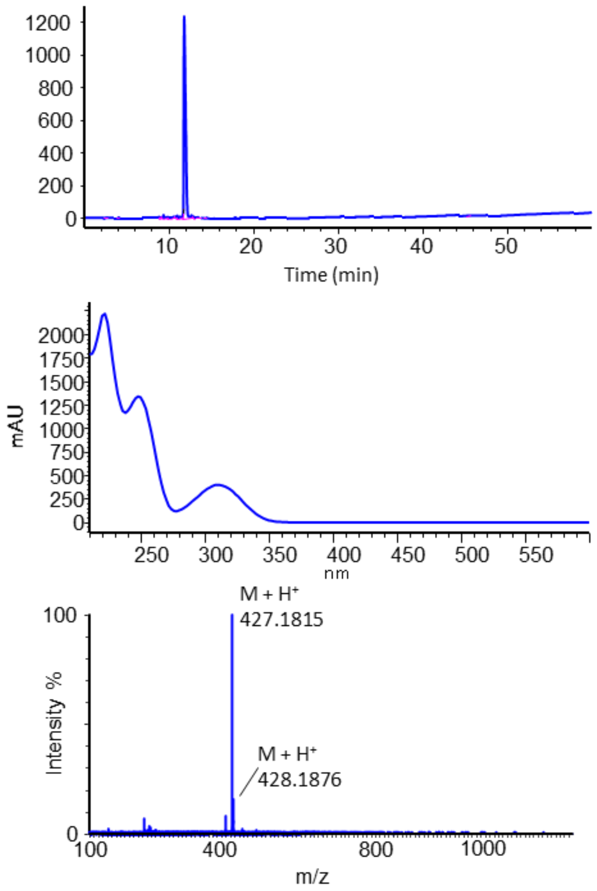


**Supplementary Figure 2** (A) HPLC trace of purified mirubactin B. (B) UV-Vis spectrum of mirubactin B. (C) HRMS spectrum showing molecular ion $[M+H]^+$ = 427.1815.

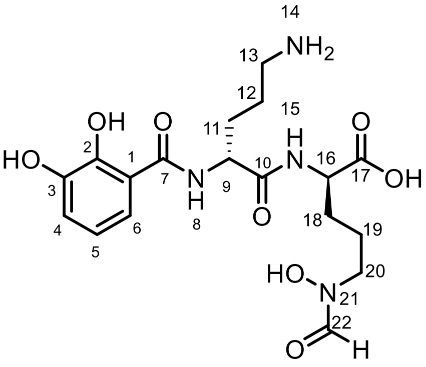


**Supplementary Table 3** NMR data for mirubactin B in $D_2O$ at 298 K, 700 MHz, $^1$H NMR collected at 700 MHz, $^{13}$C{$^1$H} NMR collected at 176 MHz, $^{15}$N shifts were measured indirectly by $^1$H-$^{15}$N HMBC performed at both 8 and 12 Hz, referencing against nitromethane

| Position | $\delta_{^1H}$ multi, (*J* in Hz) | $\delta_{^{13}C}$ | $\delta_{^{15}N}$ | HMBC |
|---|---|---|---|---|
| 1 | - | 116.4 | | |
| 2 | - | 146.3 or | | |
| 3 | - | 144.4 | | |
| 4 | 6.98 (dd, *J* = 7.9, 1.5 Hz, 1H) | 119.5 | | H to C1, 2, 3, 5, 6 |
| 5 | 6.76 (t, *J* = 8.0 Hz, 1H) | 119.6 | | H to C1, 2, 3, 4, 6, 7 |
| 6 | 7.18 (dd, *J* = 8.1, 1.6 Hz, 1H) | 119.2 | | H to C1, 2, 3, 4, 5, 7 |
| 7 | - | 169.8 | | |
| 8 | - | - | -260.5 | |
| 9 | 4.53 (dd, *J* = 7.8, 6.4 Hz, 1H) | 53.2 | | H to C1, 7, 10, 11, 12 |
| 10 | - | 172.6 | | - |
| 11 | 1.98 – 1.91 (m, 1H)<br>1.89 – 1.82 (m, 1H) | 27.8 | | H to C9, 10, 12, 13 |
| 12 | 1.80 – 1.71 (m, 2H) [overlap C18] | 23.1 | | * |
| 13 | 2.99 (ddd, *J* = 8.6, 6.9, 2.1 Hz, 2H) | 38.8 | | H to C11, 12 |
| 14 | - | - | -348.8 | |
| 15 | - | - | -252.9 | |
| 16 | 4.21 – 4.13 (m, 1H) | 54.6 | | H to C10, 17 18, 19 |
| 17 | - | 178.1 | | |
| 18 | 1.80 – 1.71 (m, 1H) [overlap C12]<br>1.69 – 1.60 (m, 1H) [overlap C19] | 28.2 | | * |
| 19 | 1.69 – 1.60 (m, 2H) [overlap C18] | 22.7 | | * |
| 20 | 3.56 – 3.44 (m, 2H) | 50.0 | | H to C16, 18, 19, 22 |
| 21 | - | - | -199.3 | |
| 22 | 7.83 and 8.18 (2 x s, 1H) | 159.4 | | H to C20 |

* Unknown due to signal overlaps

All reagents were purchased from TCI UK Ltd ($N^{\alpha}$-(5-fluoro-2,4-dinitrophenyl)-L-leucinamide, $N^{\alpha}$-(5-fluoro-2,4-dinitrophenyl)-D-leucinamide), Alfa Aesar Ltd (L-ornithine hydrochloride, 57% hydriodic acid) and Fluorochem Ltd (D-ornithine hydrochloride). LCMS was performed on a Waters LCMS system, ACQUITY Ultra Performance LC™ (UPLC™) coupled to ACQUITY Photodiode Array (PDA) UV detector and a Xevo TQ-S mass detector.

**LCMS Method for Marfey's Analysis**

HPLC separation was carried out using a linear gradient of 5 – 70 % MeCN:$H_2O$ (0.1 % formic acid) over 15 minutes and a hold time at 100% of MeCN (0.1 % formic acid) for 1 minute, with an injection volume 1 µL, and a volume flow rate of 0.25 mL/min. Detection was carried out by UV (200-500 nm) and ESI-MS. Column (temp): ACQUITY UPLC® BEH $C_{18}$ 1.7 µm, 2.1 x 150 mm Column (40 °C)

**Marfey's Analysis of mirubactin B**

To determine the absolute stereochemistry of mirubactin B by Marfey's analysis, 500 µg of mirubactin B was dissolved in 400 µL of 57 % HI and the reaction mixture heated to 110 °C in an oil bath for 3 hours. After this time the reaction mixture was evaporated to dryness under a gentle stream of nitrogen gas overnight.

The residue material was dissolved in 240 µL of 1 M aq. $NaHCO_3$, to which was added 170 µL 1 % $N^{\alpha}$-(5-fluoro-2,4-dinitrophenyl)-L-leucinamide in acetone and the reaction mixture stirred for 1 hour at 40 °C. After which the reaction was quenched with 60 µL of 2 M aq. HCl.

20 µL of this sample was diluted with 980 µL of HPLC grade water and analysed following the above LCMS method, verses standards.

**Preparation of Standards for Marfey's Analysis**

Marfey's standards for D-ornithine and L-ornithine were prepared by mixing 50 µL of a 50 mM aq. solution of either D-ornithine or L-ornithine hydrochloride with 20 µL of a 1 M aq. solution of $NaHCO_3$ and 100 µL of 1 % L-FDLA in acetone. The reaction mixtures were heated to 40 °C in a water bath for 1 hour to form the corresponding L-FDLA/D-ornithine and L-FDLA/L-ornithine adducts. 17 µL of each of these samples was diluted with 983 µL of HPLC grade water and analysed following the above LCMS method.

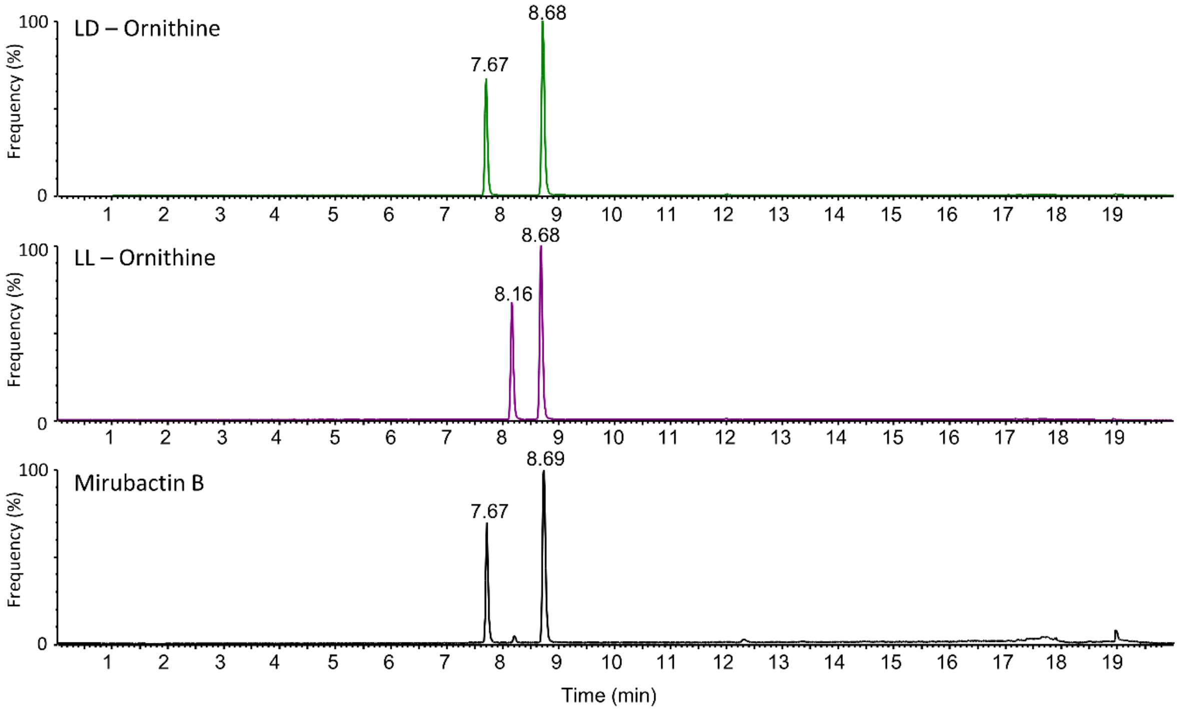


**Supplementary Figure 3** LCMS extracted ion chromatogram (EIC m/z = 427.2, +/- 0.25 Da) showing the samples of $N^{\alpha}$-(5-fluoro-2,4-dinitrophenyl)-L-leucinamide (L-FDLA) derivatized mirubactin B (bottom) in comparison to L-FDLA/D-Ornithine (top) and L-FDLA/L-Ornithine standards (middle). LCMS Extracted ion chromatogram showing the sample of L-FDLA + mirubactin B (bottom) matched with L-FDLA + D-Ornithine standard (top)

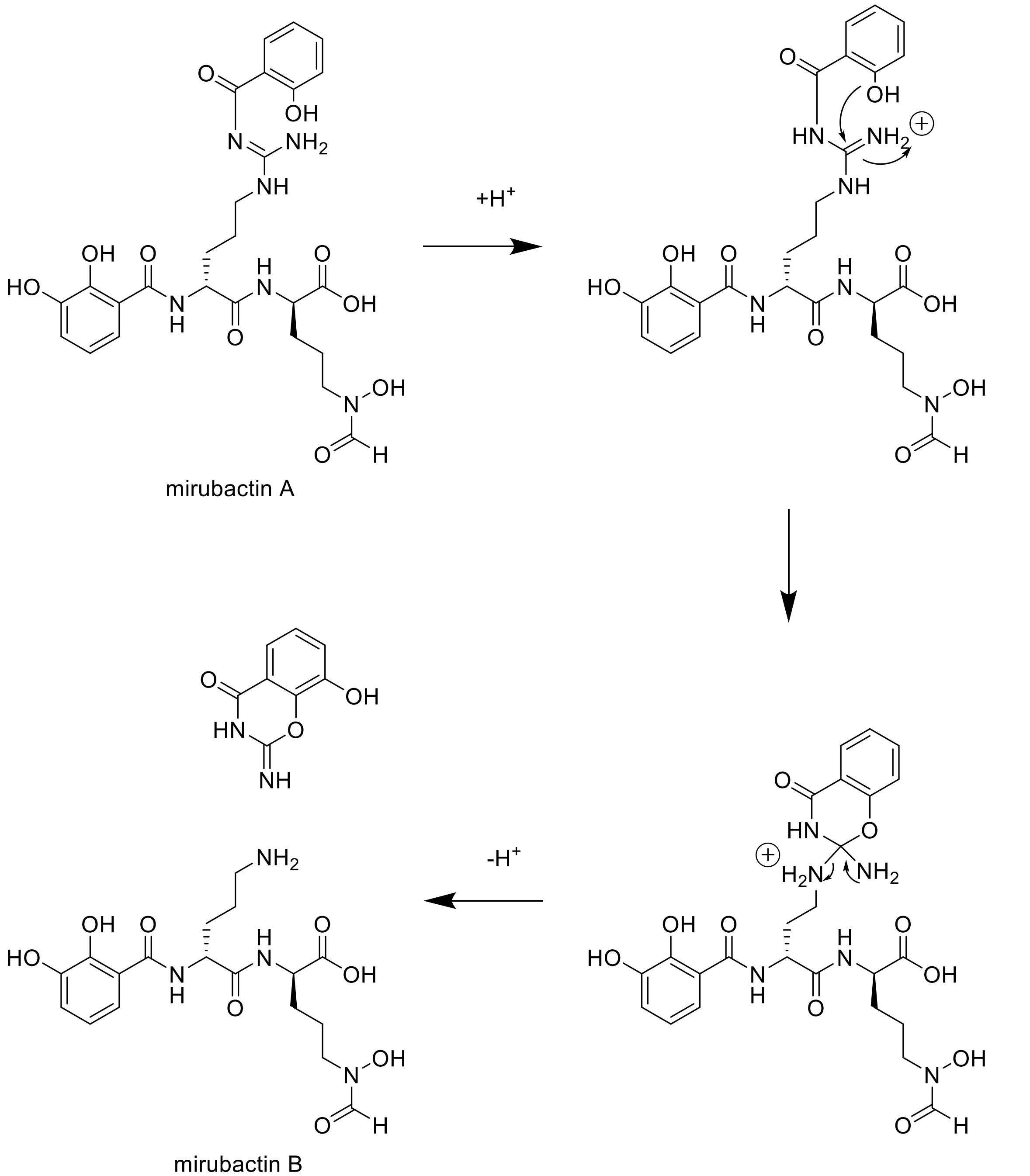


**Supplementary Figure 4** Proposed degradation pathway of mirubactin A to mirubactin B based on Kishimoto et al (2014).[34]

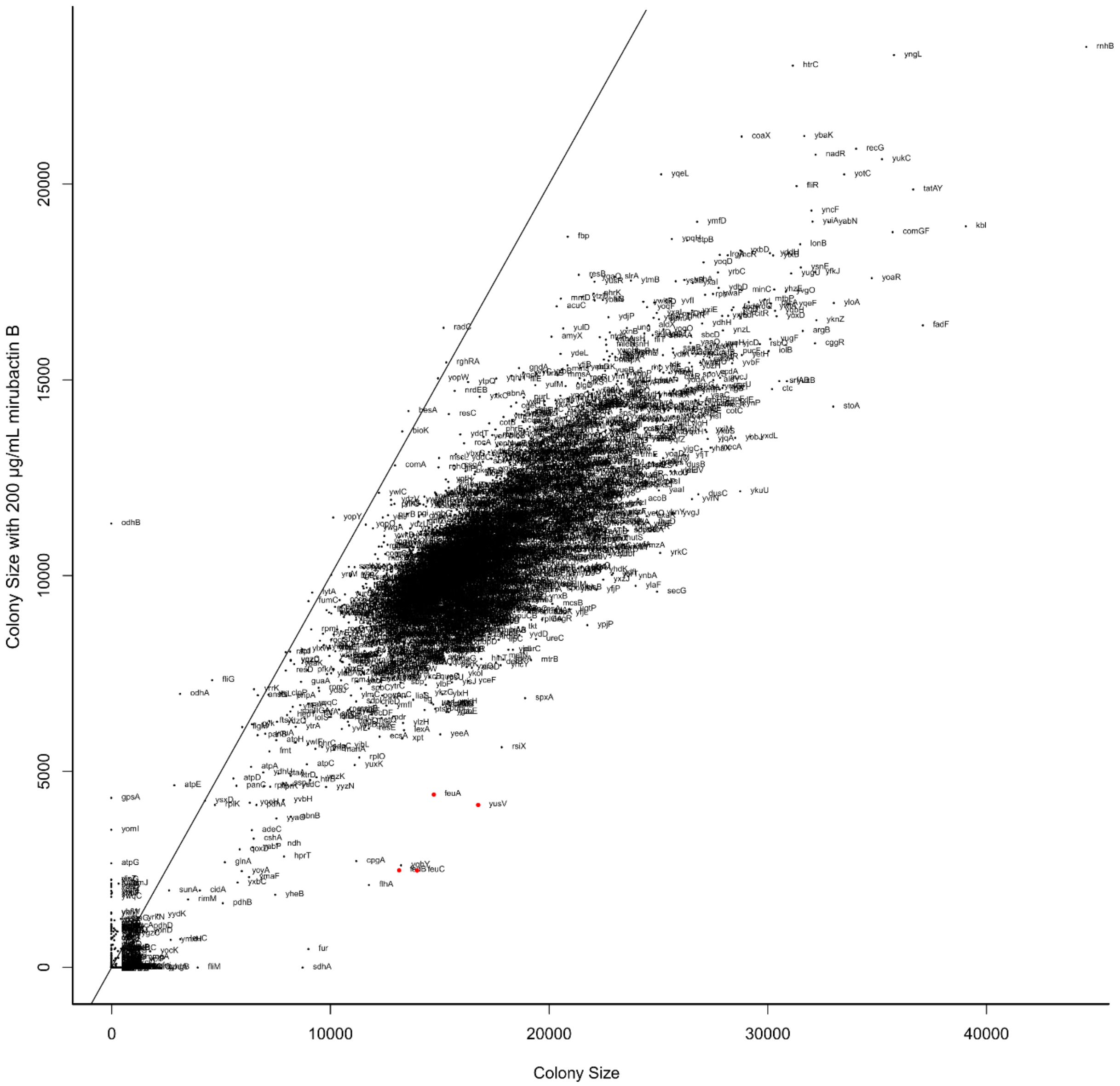


**Supplementary Figure 5** Effects of mirubactin B on the *B. subtilis* global mutant collection.

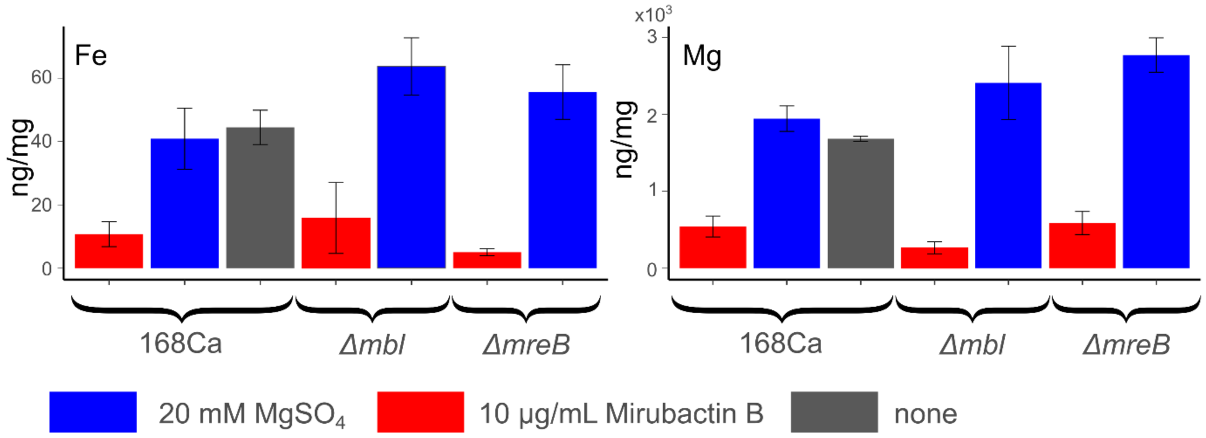


**Supplementary Figure 6** Cellular Fe and Mg content of *B. subtilis* cells, as determined by ICP-MS, after culture in PAB medium with/without supplementation with 20 mM $MgSO_4$ or 10 μg/mL mirubactin B.
